# Inducible epithelial resistance improves survival of Sendai virus pneumonia in mice by both inactivating virus and preventing CD8^+^ T cell-mediated immunopathology

**DOI:** 10.1101/2020.01.30.917195

**Authors:** S. Wali, J. R. Flores, A.M. Jaramillo, D. L. Goldblatt, J. Pantaleón García, M. J. Tuvim, B. F. Dickey, S. E. Evans

**Affiliations:** MD Anderson UTHealth Graduate School of Biomedical Sciences, Houston Texas 77030.; University of Texas MD Anderson Cancer Center, Department of Pulmonary Medicine, Houston Texas 77030.

## Abstract

Viral pneumonias remain a global health threat necessitating novel strategies to prevent and treat these lower respiratory tract infections. We have reported that mice treated with a combination of inhaled Toll-like receptor (TLR) 2/6 and TLR 9 agonists (Pam2-ODN) are broadly protected against respiratory pathogens. Although a single inhalation of Pam-ODN prevents acute morbidity and chronic complications associated with viral pneumonias, the mechanisms underlying this protection remain incompletely elucidated. Here, we show in a lethal paramyxovirus model that Pam2-ODN-enhanced survival is associated with robust virus inactivation that occurs prior to internalization by lung epithelial cells. However, it was also noted that viral mortality in sham-treated mice temporally corresponded with CD8^+^ T cell-enriched lung inflammation that peaks after the viral burden wanes. Pam2-ODN treatment also blocked this injurious inflammation, but the attenuation of lymphocytic inflammation and the reduction in virus burden were both lost when inducible reactive oxygen species generation was inhibited. Depleting CD8^+^ T cells before or after viral challenge underscored the balanced roles of CD8^+^ T cells in antiviral immunity and fatal immunopathology, but did not obviate the Pam2-ODN antiviral protection. These findings identify multifunctional inducible antiviral mechanisms and may reveal means to protect susceptible individuals against respiratory infections.

## Introduction

Viruses are the most frequent cause of community acquired pneumonia in children and adults^1^. Respiratory viral infections result in significant morbidity and mortality in vulnerable subjects, exerting a tremendous health care burden^1–4^. In addition to causing acute disease, respiratory virus infections are often complicated by chronic lung pathologies, such as asthma induction, progression and exacerbation^5–7^. Therefore, development of novel therapeutic anti-viral strategies is required to effectively prevent and treat respiratory infections and their associated chronic complications^8–10^.

While lung epithelial cells are the principal targets of most respiratory viruses, there is expanding evidence that lung epithelia are capable of generating anti-microbial responses^7,11^. We hypothesized that lung epithelial cells can be harnessed to control virus replication, thereby enhancing acute survival and reducing chronic complications of virus infections^12–15^. Our group has previously described the phenomenon of inducible epithelial resistance wherein the lungs’ mucosal defenses can be broadly stimulated to protect against a wide range of respiratory pathogens, including viruses^12–17^. This protection is induced by a single inhalation of a combination treatment consisting of Toll like receptor (TLR) 2/6 and 9 agonists (Pam2-ODN) before or after viral challenge. While no individual leukocyte populations have been identified as critical for Pam2-ODN-induced resistance, lung epithelial cells are essential to the inducible anti-viral response^12^. Further, we have shown that Pam2-ODN mediated protection is dependent upon epithelial generation of reactive oxygen species (ROS) but, interestingly, does not require Type I interferons^16,17^. More recently, we have shown prevention of chronic virus-induced asthma in mice treated with Pam2-ODN but we have not clarified the anti-viral mechanisms^18^.

In this study, we investigated the mechanisms of Pam2-ODN enhanced mouse survival of paramyxovirus, Sendai virus (SeV) infection. We found that Pam2-ODN treatment not only reduced lung SeV burden but also decreased epithelial cell injury and host immunopathologic leukocyte responses to SeV infections. While CD8^+^ T cells are known to contribute to virus clearance, it is shown here that CD8^+^ T cells also cause substantial mortality that can be prevented by Pam2-ODN treatment. Notably, Pam2-ODN-induced enhanced survival of SeV infection is effective even in CD8^+^ T cell deficient conditions. Further, we demonstrate anti-viral mechanisms of inducible epithelial resistance, where virus particles are inactivated in a ROS-dependent manner prior to internalization by their epithelial targets.

## Results

### Enhanced mouse survival of SeV infection by Pam2-ODN treatment

Aerosolized Pam2-ODN treatment one day prior to SeV challenge increased mouse survival of SeV challenge and reduced mouse weight loss (Fig. 1a, b), similar to the protection observed against lethal influenza pneumonia^12,15,16^. The survival benefit was associated with reduced lung SeV burden, as measured by SeV M gene expression (Fig. 1c). Investigating the natural progression of infection revealed that SeV lung burden was maximal on day 5 and gradually decreased until falling below the limit of quantification (LOQ) by day 11 (Fig. 1d). Pam2-ODN pretreatment reduced SeV burden on all assessed days (Fig. 1d). Although the lethality of SeV infection was highly dependent on the inoculum size, we found that peak mortality paradoxically occurred around days 10 to 12 irrespective of inoculum size, despite the fact that SeV is essentially undetectable that long after challenge (Fig. 1a, d, e). Assessing the effect of Pam2-ODN on SeV burden in immortalized mouse epithelial cells (MLE-15) and primary mouse tracheal epithelial cells (mTEC), we found that Pam2-ODN treatment reduced SeV burden at every time point measured, reflecting the inducible antiviral capacity of isolated epithelial cells (Fig. 1f).

**Figure 1.**
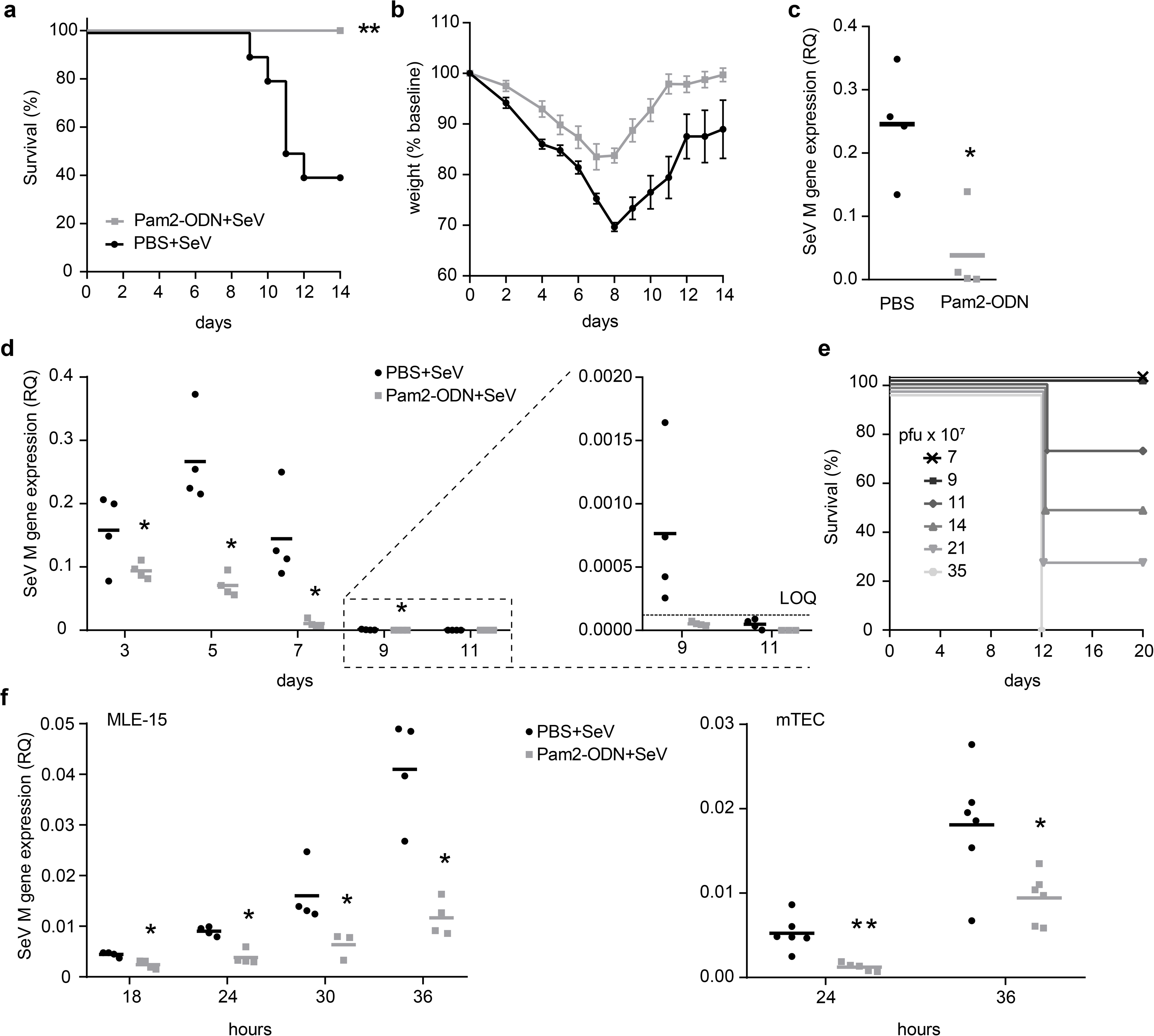
Pam2-ODN enhances mouse survival of SeV infection and reduces lung virus burden. Survival **(a)** and weight loss **(b)** of mice treated with PBS or Pam2-ODN one day prior to SeV virus challenge. **(c)** Mouse lung SeV burden 5 days after infection assessed by qPCR for Sendai Matrix (M) gene (Relative quantification, RQ to 18S) relative to 18S. **(d)** Time course of lung SeV burden in mice treated with PBS or Pam2-ODN; LOQ, limit of quantification. **(e)** SeV inoculum dependent mouse survival. **(f)** SeV burden assessed by qPCR in MLE-15 cells and primary mouse tracheal epithelial cells (mTEC) treated with PBS or Pam2-ODN 4h prior to SeV challenge. n=10 mice per group in survival plots, n=4 mice/group in virus burden experiments. \**p*<0.05, \*\**p*<0.005.

### Pam2-ODN treatment attenuates SeV-induced epithelial injury

This temporal dissociation between peak virus burden and peak mortality led to the hypothesis that SeV-induced mortality may not be exclusively driven by excessive virus burden but may also result from untoward SeV-induced host immune response. Therefore, the acute changes in mouse lungs following SeV infection were characterized. We found increases in lung epithelial cleaved caspase 3 (cCasp3), a marker for programmed cell death, on days 7 to 11 after SeV infection (Fig. 2a, upper panel). Virus infection-related epithelial cell injury and death is typically associated with proliferative repair mechanisms^19,20^. Staining the infected mouse lung tissue for Ki67 and EdU revealed maximum signals for both markers in the second week after infection (Fig. 2b-e, upper panel). These events of lung epithelial cell death and proliferation coincided with the peak of mortality (day 12, Fig. 1e). Further, hematoxylin and eosin staining of lung tissues infected with SeV showed profound increases in inflammatory cells from days 7 to 10 with evidence of damaged airway and parenchymal tissue (Fig. 2f). However, Pam2-ODN pretreatment of mice reduced epithelial cell injury and proliferation (Fig. 2a-e, lower panel). This temporal association of epithelial injury and death after viral clearance supported our hypothesis that mouse mortality caused by SeV infection is due in part to the host immune response to SeV infections.

**Figure 2.**
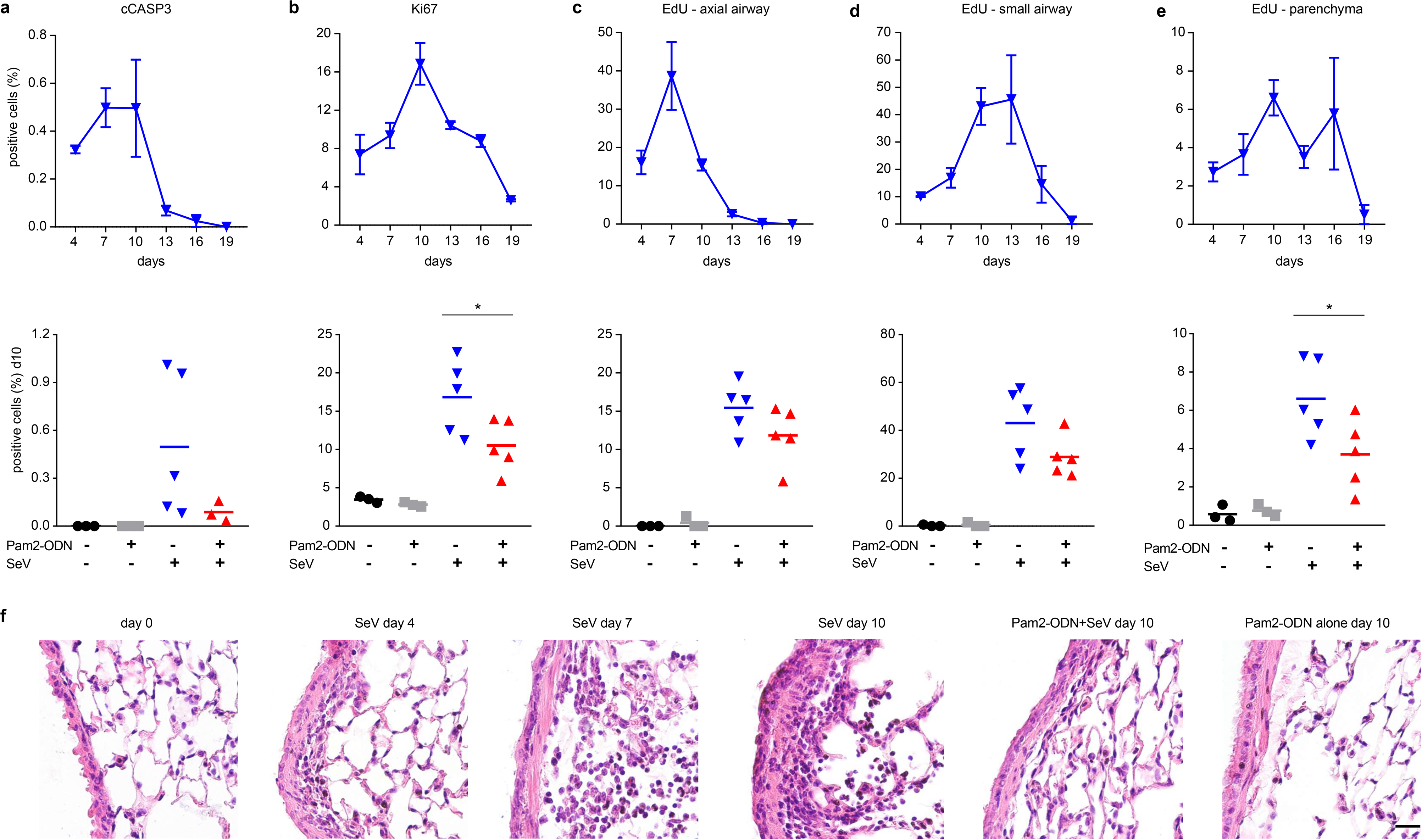
Pam2-ODN pretreatment reduces epithelial cell death and proliferation during acute SeV infection. Cleaved caspase 3 **(a)** or Ki67 **(b)** positive cells in mouse lung epithelium after SeV infection with or without Pam2-ODN treatment (lower panel). EdU positive cells in axial **(c)**, small airways **(d)** and parenchyma **(e)** after SeV infection with or without Pam2-ODN (lower panel). **(f)** Mouse lung histology following SeV challenge with or without Pam2-ODN. n=5 mice per condition. Scale bar = 100 μm. \**p*<0.05.

### Pam2-ODN attenuates SeV-induced lymphocytic lung inflammation

To explore this hypothesis, the host leukocyte response to SeV infection was characterized. Differential Giemsa staining of bronchoalveolar lavage (BAL) cells revealed increased neutrophils on days 2 to 5 and increased macrophages on days 5 to 8 (Fig. 3a, left and middle panel, solid grey line) after SeV challenge. Congruent with our prior studies, inhaled treatment with Pam2-ODN in the absence of infection led to a rapid rise in neutrophils that was resolved within 5 days (Fig 3a, dashed line)^21^. The neutrophil response to SeV challenge was modestly increased among mice pretreated with Pam2-ODN (Fig. 3a, left panel, solid dark line). Pam2-ODN-treated, SeV-challenged mice showed almost no difference in macrophages compared to PBS-treated, SeV-challenged mice (Fig. 3a, middle panel, solid dark line). A rise in lymphocytes was observed on days 8 to 11 in PBS-treated, SeV-challenged mice (Fig. 3a, right panel, solid grey line), temporally corresponding with peak mortality. However, Pam2-ODN treated, SeV-challenged mice displayed significantly reduced lymphocyte numbers at every time point assessed (Fig 3a, right panel, solid dark line). The gating strategy for lymphocyte subsets by flow cytometry is shown in Supplementary Fig. 1. A modest reduction in CD4^+^ T cells was observed in Pam2-ODN-treated, SeV-challenged mice compared to PBS-treated, SeV-challenged mice (Supplementary Fig. 2). We also found the percentage of CD19^+^ B220^+^ B cells reduced after SeV infection in comparison to Pam2-ODN treated and uninfected mice (Supplementary Fig. 2), as has been seen with other viral models^22,23^. However, the biggest difference between groups was in CD8^+^ T cells, with Pam2-ODN-treated, SeV-challenged mice displaying a significantly lower number and percentage of CD8^+^ T cells than PBS treated, SeV-challenged mice (Fig. 3b, c). Since the greatest difference after Pam2-ODN treatment was in CD8^+^ T cell levels and there was a tight correlation between peak mortality and the increase in lung CD8^+^ T cells on days 8 to 11, we investigated the role of CD8^+^ T cells in SeV-induced mortality.

**Figure 3.**
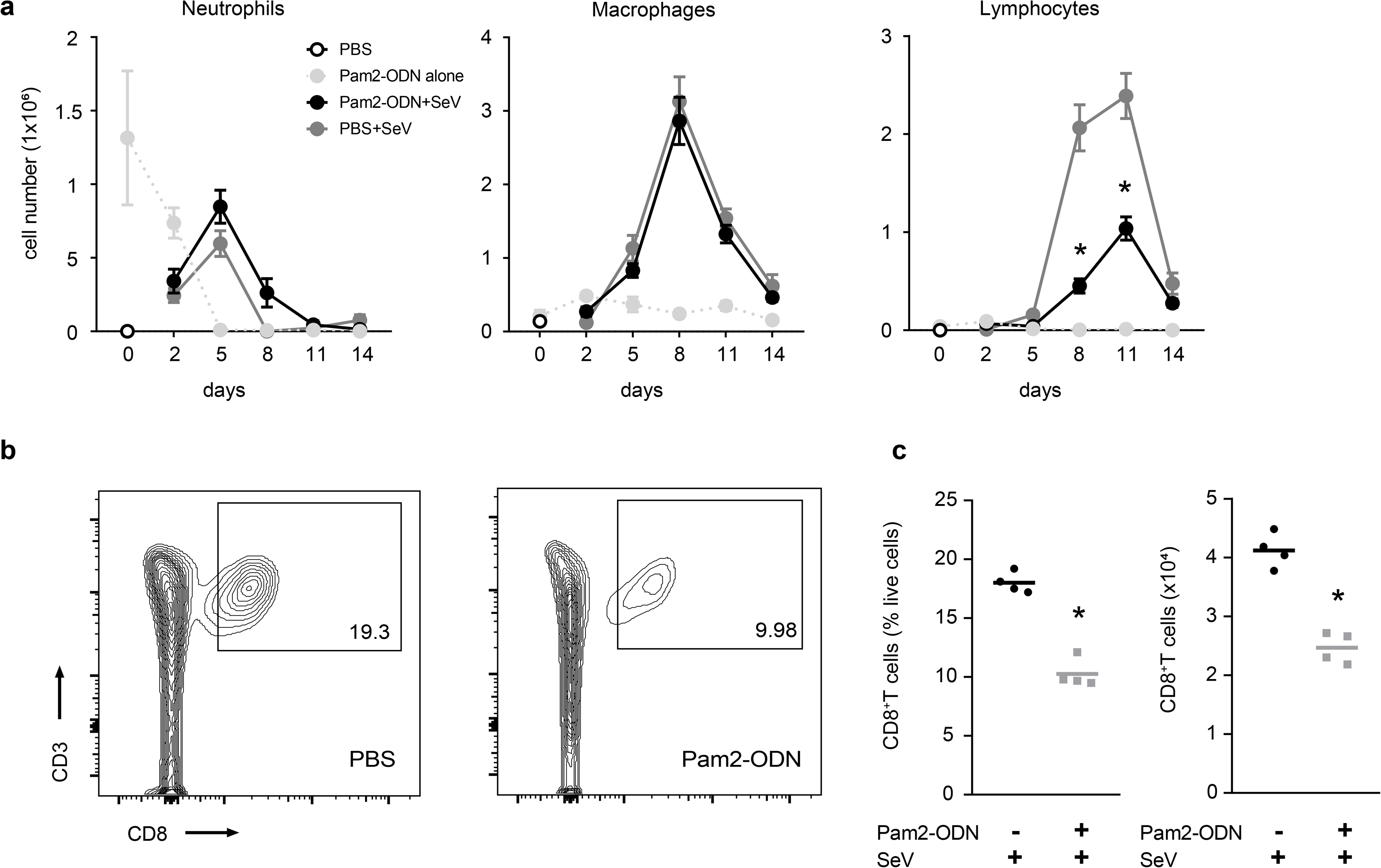
Pam2-ODN pretreatment reduces SeV induced CD8^+^ T cells. **(a)** Differential Giemsa staining of BAL cells from mice challenged with SeV with or without Pam2-ODN pretreatment. **(b)** Flow cytometry for CD8^+^ T cells following SeV infection with or without Pam2-ODN. **(c)** Lung CD8^+^ T cells 11 days after SeV challenge in mice pretreated with PBS or Pam2-ODN. \**p*<0.05

### CD8^+^ T cells contribute to anti-viral immunity but also cause lethal immunopathology

To assess the role of CD8^+^ T cells in SeV defense and host mortality, mouse CD8^+^ T cells were depleted prior to SeV challenge (Fig. 4a, Supplementary Fig. 3). This depletion resulted in significantly reduced survival of SeV infection (Fig. 4b). The nearly 90% mortality in SeV challenged CD8^+^ T cell depleted mice (Fig. 4b) was associated with impaired viral clearance (Fig. 4c, Supplementary Fig. 4). This was not surprising, given the known role of CD8^+^ T cells in virus clearance^24–28^. However, Pam2-ODN treatment still significantly enhanced survival of SeV challenge, even in the absence of CD8^+^ T cells (Fig. 4b). This protection was again associated with reduced SeV burden compared to CD8^+^ T cell-depleted, SeV-challenged mice without Pam2-ODN treatment (Fig. 4c, Supplementary Fig. 4). These results were congruent with our previous studies showing Pam2-ODN inducible resistance against bacterial pneumonia despite the lack of mature lymphocytes (*Rag1*^−/−^)^12^.

**Figure 4.**
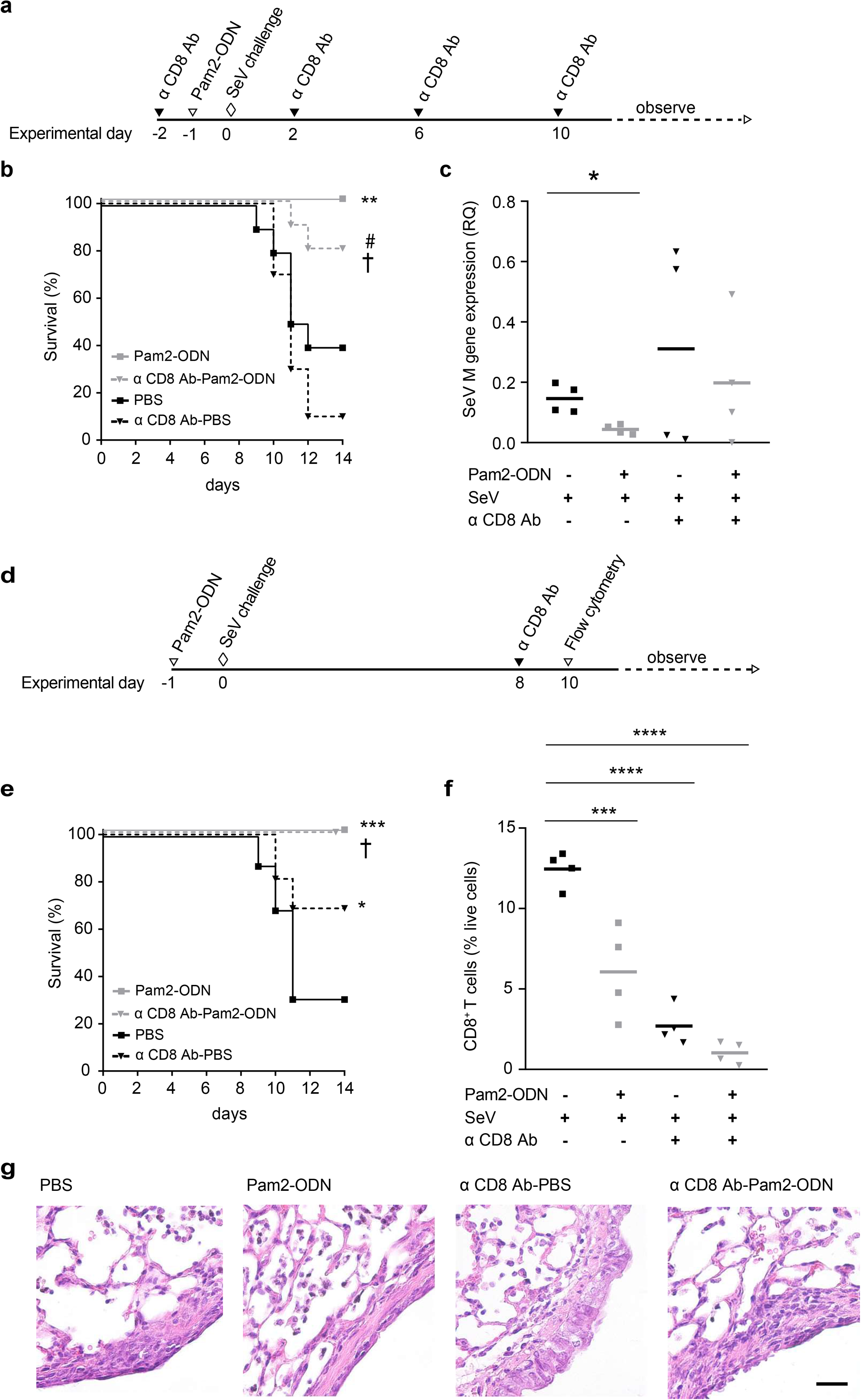
Pam2-ODN treatment reduces CD8^+^ T cell mediated SeV induced immunopathology. Experimental outline **(a)**, survival **(b)** and SeV burden **(c)** 5 days after SeV challenge following PBS or Pam2-ODN treatment and with or without preinfection CD8^+^ T cell depletion. Experimental outline **(d)**, survival **(e)** and CD8^+^ T cells **(f)** 10 days after SeV challenge following pretreatment with PBS or Pam2-ODN and with or without CD8^+^ T cells depleted on day 8 of SeV challenge. **(g)** Lung histology 10 days after SeV challenged with or without Pam2-ODN treatment and/or CD8^+^ T cells. Scale bar = 100 μm. n=10 mice/group for survival in experiment **a** and **b**; n=16 mice/group for survival in experiment **d** and **e**. \*\*\*\**p*<0.0001 compared to PBS, \*\*\**p*<0.0005 compared to PBS, \*\**p*<0.005 compared to PBS, #*p*<0.005 compared to α CD8 Ab-PBS, †*p*<0.05 compared to PBS, \**p*<0.05 compared to PBS, \**p*<0.05 compared to α CD8 Ab-PBS, †*p*<0.05 compared to α CD8 Ab-PBS.

As it appeared likely that CD8^+^ T cells contributed both beneficial (antiviral) and deleterious (immunopathologic) effects, we depleted the CD8^+^ T cells on day 8 -- after virus begin to clear but before peak mortality (Fig. 4d). Interestingly, mice depleted of CD8^+^ T cells on day 8 displayed enhanced survival of SeV challenge compared to mice with intact CD8^+^ T cells (Fig. 4e). Depletion of CD8^+^ T cells was confirmed by flow cytometry in disaggregated lung cells 10 days after SeV challenge (Fig. 4f, Supplementary Fig. 3). We also assessed lung injury by H&E staining of lung tissue 10 days after SeV challenge and found increased inflammation and epithelial cell damage in undepleted mice compared to CD8^+^ T cell-depleted mice (Fig. 4g). This supported our hypothesis that, while CD8^+^ T cells confer anti-viral immunity, they also contribute to fatal SeV-induced immunopathology.

### Pam2-ODN treatment leads to extracellular inactivation of virus particles

As the antiviral protection consistently correlated with reduced viral burden, and as the reduced virus burden likely contributes to the reduced CD8^+^ T cell levels, we sought to determine how Pam2-ODN-induced responses cause antiviral effects. We investigated whether the principal Pam2-ODN effect occurred before (extracellular) or after (intracellular) virus internalization into their epithelial targets. Using multiple methods to determine the effect of Pam2-ODN on SeV attachment, we found no differences in attachment (Fig. 5a-d). However, even though similar numbers of virus particles were attached to epithelial cells, when these attached virus particles were liberated from the epithelial cell targets, virus particles from Pam2-ODN-treated epithelial cells were less able to subsequently infect other naive epithelial cells (Fig. 5e, f). As the number of attached virus particles was the same, this difference in SeV burden in cells that received liberated virus particles from PBS vs Pam2-ODN treated cells indicated that SeV is inactivated prior to epithelial internalization.

**Figure 5.**
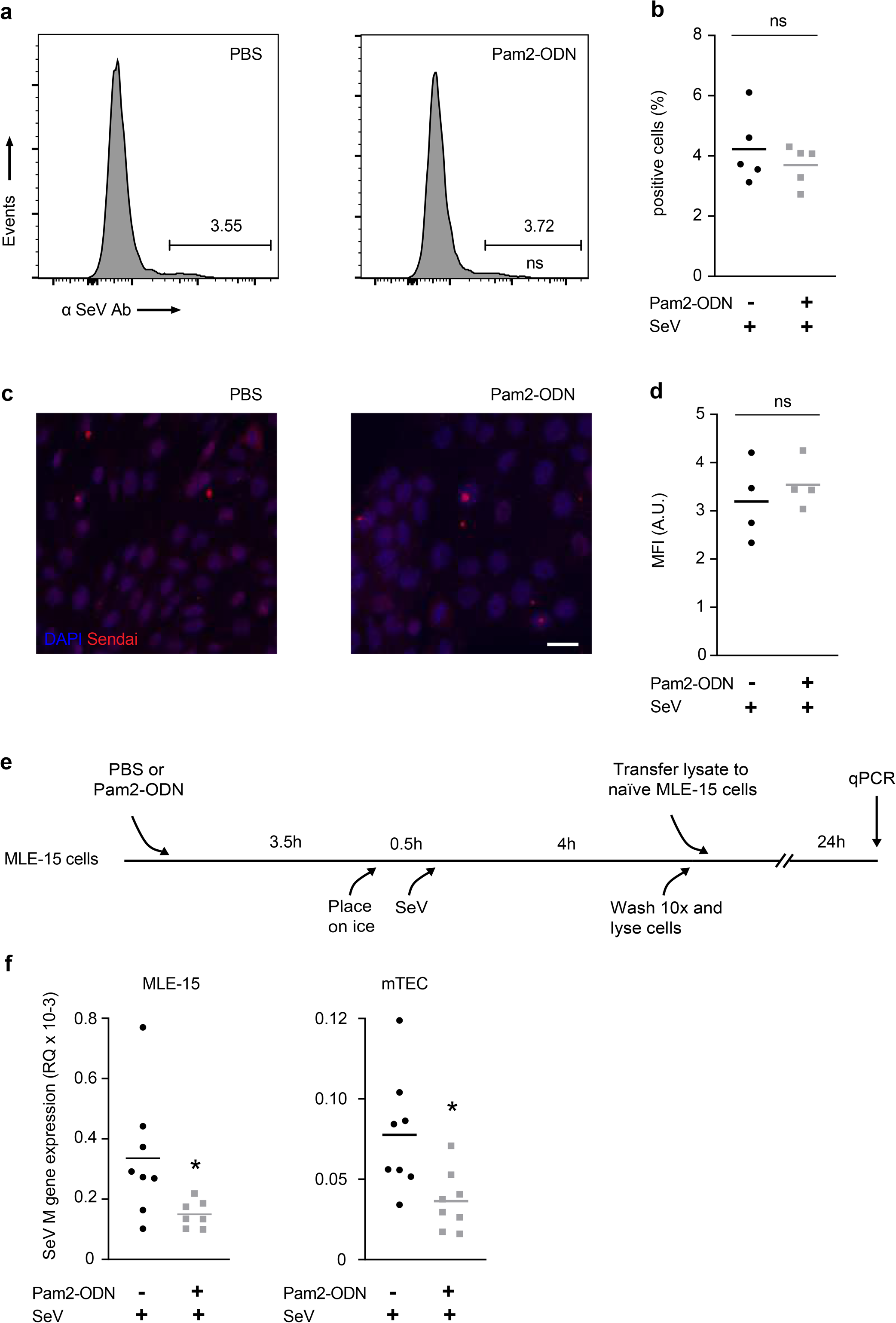
Pam2-ODN inhibits SeV without altering attachment. **(a)** Flow cytometry to measure virus attachment to epithelial cells. **(b)** Percentage of SeV positive epithelial cells from **(a)**. **(c)** Representative examples of immunofluorescence for virus attachment. **(d)** Mean fluorescence intensity of SeV-exposed epithelial cells. **(e)** Experimental outline. **(f)** SeV M gene expression in MLE-15 cells (left) or primary tracheal epithelial cells (right) receiving liberated virus from cultures pretreated with PBS or Pam2-ODN prior to SeV infection. \**p*<0.05

### Pam2-ODN-induced epithelial ROS protect against SeV infection and CD8+ T cell immunopathology

The anti-influenza response initiated by Pam2-ODN requires epithelial generation of ROS from both NADPH-dependent dual oxidase and mitochondrial sources^16,17^. Extending these findings to the SeV model, an NADPH oxidase inhibitor (GKT 137831) fully abrogated the Pam2-ODN-induced anti-SeV response (Fig. 6a). Similarly, treatment with a combination of FCCP (uncoupler of oxidative phosphorylation) and TTFA (complex II inhibitor) obviated the Pam2-ODN-induced anti-SeV response (Fig. 6b)^16,17^. Further, it was found that Pam2-ODN induced epithelial generation of ROS were required for inactivation of SeV prior to epithelial entry (Fig. 6c). Congruent with these *in vitro* and *ex vivo* studies, mice treated with FCCP-TTFA before Pam2-ODN treatment and SeV challenge (Fig. 6d) demonstrated reduced survival (Fig. 6e), increased SeV burden (Fig. 6f), and increased CD8^+^ T cells on day 10 (Fig 6g).

**Figure 6.**
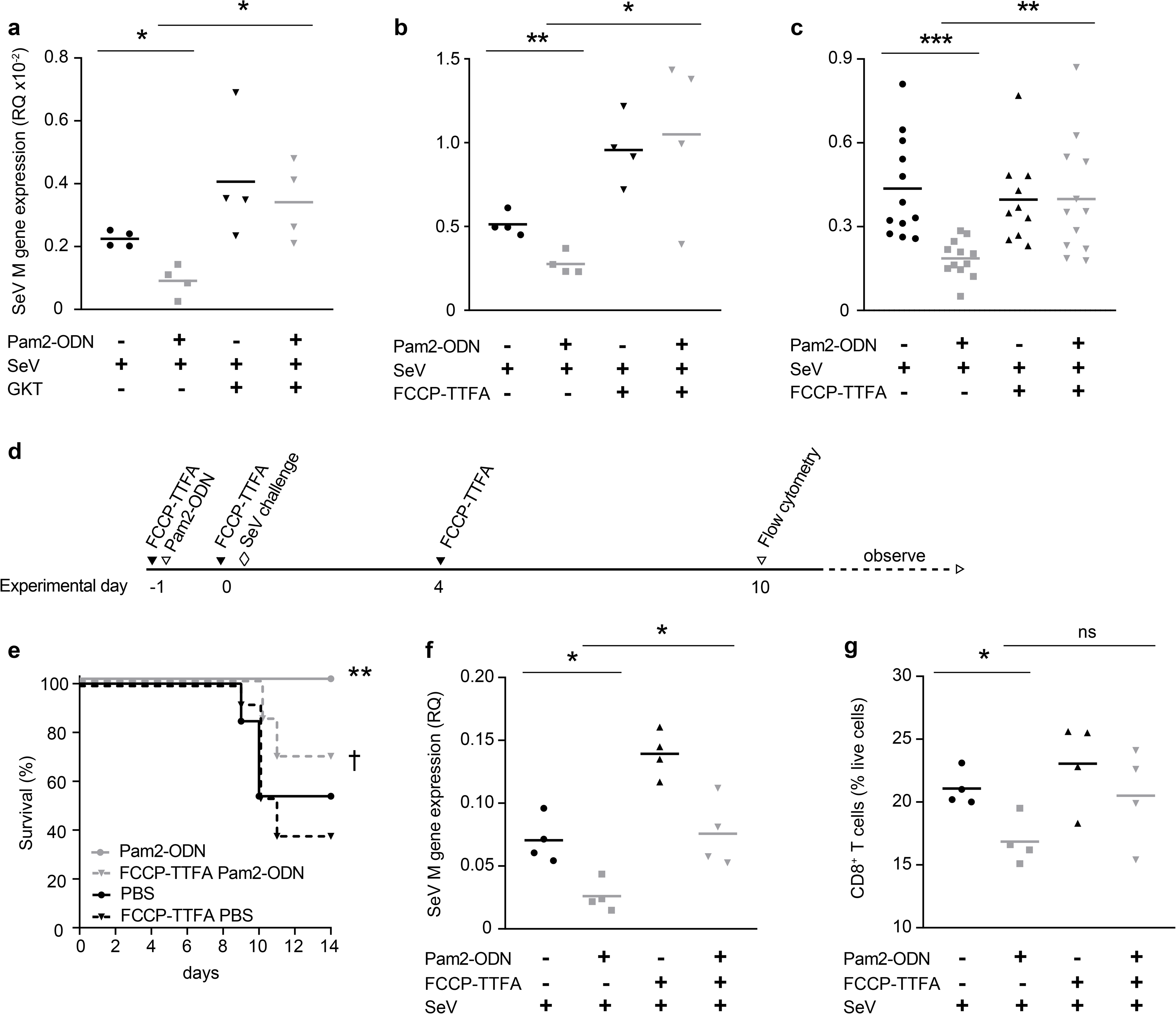
Pam2-ODN induced reactive oxygen species protects against acute SeV virus infections and immunopathology. SeV burden in MLE-15 cells with or without treatment with Pam2-ODN and/or NADPH inhibitors **(a)** or mitoROS inhibitors **(b)**. **(c)** SeV M gene expression in MLE-15 cells receiving liberated virus from cells pretreated with PBS/Pam2-ODN with or without mitoROS inhibition. **(d)** Experimental outline. **(e)** Survival of SeV challenge in mice treated with PBS or Pam2-ODN and/or mtROS inhibitors. **(f)** Lung SeV burden measured on day 5 and **(g)** lung CD8^+^ T cells assessed on day 10. n=13 mice/group in experiment **d** and **e**. \*\*\**p*<0.0001, \*\**p*<0.005, \*\**p*<0.01 compared to PBS, †*p*<0.05 compared to Pam2-ODN, \**p*<0.05, \**p*<0.01.

## Discussion

In this study, we demonstrate that therapeutic stimulation of lung epithelial cells enhances mouse survival of acute SeV infections by both reducing the virus burden and attenuating host immunopathology. While our group has demonstrated inducible resistance against multiple respiratory pathogens including viruses^12–17,21^, these studies demonstrate for the first time when in the virus lifecycle the anti-viral effects begin, substantiate the role of ROS in SeV protection, and reveal protective immunomodulatory effects.

While Pam2-ODN treatment provided a significant host survival benefit in SeV infection, we observed this survival difference occurring even after the PBS-treated mice had cleared the virus. This observation prompted the hypothesis that host mortality is not the exclusive result of direct viral injury to the lungs, but due at least in part to the host response to the virus infections. Eliminating the anti-viral component of CD8^+^ T cells in this model, we observed enhanced survival of SeV infections in mice depleted of CD8^+^ T cells on day 8 (Fig. 4), revealing the importance of balancing the dual functions of CD8^+^ T cells in anti-viral immunity and in causing fatal immunopathology. Our findings suggest that the surge in CD8^+^ T cells within the lungs after most virus has been cleared causes physiologic impairment via lung injury and cell death (Fig 4d-g). However, Pam2-ODN treatment enhanced survival of SeV infections, reflecting the intrinsic anti-viral capacity of the lung epithelial cells. This finding is potentially valuable in the context of treating pneumonia in immunocompromised patients.

Previous reports support this concept of counter-balanced immune protection and immunopathology by CD8^+^ T cells during virus infections^24,28–32^. Some reports have shown that antigen-experienced memory CD8^+^ T cells enhance respiratory syncytial virus (RSV) clearance, but also mediate severe immunopathology^30,33^. However, our study is the first to demonstrate the survival advantage of paramyxovirus infections by either temporal depletion of enhanced CD8^+^ T cells or by inducible epithelial resistance. Our findings are also congruent with reports on the role of CD8^+^ T cells in non-respiratory viral infection models, such as in West Nile virus infection, where CD8^+^ T cell deficient mice display decreased mortality^31^. Findings from this study and others reveal that the harmful effects of CD8^+^ T cell mediated immunopathology may supersede the benefits of T cell mediated viral clearance. Therefore, it is appealing to develop inducible anti-microbial strategies that do not rely on conventional T cell-mediated microbial clearance and are also effective in vulnerable immune deficient populations^12,16,34,35^.

Although the CD8^+^ T cell depletion studies enhanced our understanding of immunopathology in virus infections, much of the survival benefit against SeV infection was presumably mediated by anti-viral effects induced by Pam2-ODN. This led us to investigate the mechanisms of these inducible anti-viral effects. Given the multiple steps in the virus life cycle, it was not known at what stage Pam2-ODN exerted its anti-viral effect. Exploring this, we found no difference in number of SeV particles attached to the cells between PBS and Pam2-ODN treatment (Fig 5a-d). However, attached virus particles that were from Pam2-ODN treated cells retain less infective capacity when added to naïve epithelial cells, revealing pre-internalization virus inactivation by Pam2-ODN treatment (Fig 5e, f).

Knowing that Pam2-ODN inducible resistance required ROS production to protect against influenza^16^, we studied the role of ROS in Pam2-ODN-mediated reduction in SeV burden. ROS inhibition not only led to attenuation of Pam2-ODN’s anti-viral effect but was also permissive for increased lung CD8^+^ T cell numbers *in vivo*, implicating Pam2-ODN-induced ROS in preventing both identified mechanisms of mouse mortality in SeV pneumonia (Fig 6a, b, e, f). Further, ROS inhibition also led to loss of Pam2-ODN-inducible *in vitro* inactivation of SeV prior to epithelial internalization (Fig. 6c), demonstrating for the first time that epithelial ROS directly contribute to virus inactivation.

Production of ROS as a microbicidal mechanism has been widely reported in phagocytic cells^36–38^. However, this mechanism has not been extensively studied in non-phagocytic cells^39^, where it apparently acts predominantly extracellularly rather than intracellularly as in phagocytes. (Fig. 5f, g). Induction of a sustained microbicidal state through lung epithelial reprogramming can potentially explain the broad protection seen against multiple pathogens^12–14,21,34,40^. These findings of viral inactivation by epithelial ROS production reveal an essential component of inducible epithelial resistance.

Taken together, these findings provide mechanistic insights into the role of the lung epithelium in induction of anti-viral responses and prevention of host immunopathology that may inform future therapeutics to protect vulnerable populations.

## Methods

### Mice

All *in vivo* experiments were performed using 6 to 10-week-old C57BL/6J mice of a single sex purchased from (Jackson laboratory) or bred in-house according to the Institutional Animal Care and Use Committee of MD Anderson Cancer Center, protocol 00000907-RN01.

### Cells

Mouse lung epithelial (MLE-15) cells were kindly provided by Jeffrey Whitsett, Cincinnati Children’s Hospital Medical Center, and cultured in DMEM with 2 % Fetal Bovine Serum (FBS), 1 % insulin and transferrin. MLE-15 cells were authenticated by the MD Anderson Characterized Cell Line Core Facility. To harvest tracheal epithelial cells, mice were anesthetized to expose and excise tracheas. These tracheas were then digested in pronase (1.5 mg/ml, Sigma Aldrich) overnight at 4° C. Tracheal epithelial cells were then isolated and cultured on collagen coated transwells in Ham’s F12 media supplemented with differentiation growth factors and hormones as described previously^16,35^.

### TLR treatments and viral challenge

For *in vitro* treatments, cells were treated with Pam2 (2.2 μM) and ODN (0.55 μM), 4 h before SeV inoculation as previously described^16,17^. For *in vivo* treatments, 10 ml solution of Pam2 (4 μM) and ODN (1 μM) in endotoxin free water was delivered by Aerotech II nebulizer (Biodex Medical Systems) driven by 10 l/min along CO2 (5 %) in air for 30 minutes as previously described^16,17^. SeV was purchased from ATCC (Manassas, VA) and grown in Rhesus monkey cells obtained from Cell Pro labs (Golden Valley, MN). For *in vitro* inoculations, multiplicity of infection (MOI) of one was used. Unless otherwise stated, mice were challenged with 1 × 10^8^ plaque forming units (pfu) in PBS inserted into the oropharynx of mice, under isoflurane anesthesia as described^18^. Mice were weighed before and daily after challenge as a measure of morbidity and criteria for euthanasia.

### Bronchoalveolar lavage and differential Giemsa staining

Mouse tracheas were instilled with 1.5 ml of PBS through a 20-gauge cannula after deep sedation and approximately 1 ml of BAL fluid was collected. The BAL fluid was then spun down at 4° C at 300 *g* to collect the cells in the pellet. The cell pellet was resuspended in 1 ml of ice-cold PBS and 200 μl of this cell suspension was then subjected to cytocentrifugation at 300 *g* for 5 min. Cells were stained with Giemsa stain for differential counts determination and total cells were counted by hemocytometer.

### Flow cytometry

For *in vivo* experiments, mouse lungs were perfused with 5 to 10 ml PBS, dissected, cut into 1 mm^3^ pieces, and digested with collagenase/DNAse I (5 mg/ml, Worthington biochemical) for 30 min at 37° C. After digestion, single cells were collected by passing through a 70 μm filter. These single cells were washed with FACS staining buffer (PBS supplemented with 1 % FBS) and stained for specific cell types, as indicated in the antibody table (Table 1). For *in vitro* experiments, MLE-15 cells were seeded on 24 well plates for treatment with Pam2-ODN and SeV inoculation. Cells were trypsinized and washed with FACS staining buffer 2X. Cells were blocked in 5 % donkey serum for 30 min before proceeding to staining with Rabbit SeV antibody (MBL International) overnight at 4° C, followed by staining with secondary Alexa488 anti-rabbit antibody (Jackson Immunologicals) for 1 h. Cells were fixed and acquired on BD LSRII (BD Biosciences) for Alexa488 positive cells.

**Table 1.**

| **Antibodies** | **Vendor** | **Catalogue numbers** |
| --- | --- | --- |
| CD3 | Tonbo | 65-0031-U100 |
| CD4 | Tonbo | 60-0042-U100 |
| CD8 | Tonbo | 25-0081-U100 |
| Live dead | Tonbo | 13-0870-T500 |
| CD25 | Biolegend | 102038 |
| Foxp3 Treg kit | eBiosciences | 72-5775 |
| CD8-Depleting<br>Ab | Bioxell | BE0223-A025 |
| CD19 | Biolegend | 115507 |
| B220 | BD Biosciences | 562922 |
| Anti-SeV virus<br>Ab | MBL International | PD029 |
| Ki67 | Invitrogen | MA5-14520 |
| cCasp3 | Cell signaling | 9662S |

### Immunofluorescence

MLE-15 cells were grown on chamber slides (Labtek), treated with Pam2-ODN for 4 h before inoculation with SeV (MOI 1). Cells were then fixed with 2 % paraformaldehyde before staining with Rabbit SeV antibody (MBL International) and detected using a secondary anti-rabbit antibody. For each experimental condition, specimens were imaged using Olympus BX 60 microscope using identical parameters for time of exposure, color intensity, contrast and magnification. Images were then loaded on ImageJ software to calculate mean fluorescence intensity for each group.

### H&E staining

Mouse lungs were fixed by intratracheal inflation with 10 % formalin for 24 h, and then transferred to 70 % ethanol embedded in paraffin. Tissue blocks were then cut into 5 μm sections, mounted onto frosted glass slides, deparaffinized with xylene, washed with ethanol, then rehydrated and stained with hematoxylin and eosin for morphological changes.

### Epithelial proliferation assays

Mice were injected intraperitoneally with 0.1 ml EdU (1 mg/mouse). After 24 h, lungs were inflated and fixed with 10 % formalin for 24 h at 4° C and then lungs were embedded in paraffin. Paraffin sections were cut into 5 μm transverse sections of the axial airway, between lateral branches 1 and 2. Lung sections were then stained following the Click-iT EdU Imaging Kit protocol for EdU (Abcam,) followed by staining with DAPI for 30 min at room temperature. Images were collected using Olympus BX 60 microscope using identical parameters for all conditions. Some lung sections were subjected to antigen retrieval and then stained for Ki67 (1:1000; Invitrogen) and cCasp3 (1:500; Cell Signaling). EdU, Ki67 or cCasp3 positive cells were quantified using cell counter plugin in ImageJ and normalized to DAPI positive cells in every field of view (number of fields surveyed per mouse sample = 3).

### CD8^+^ T cell depletion

Anti CD8-β antibody (200 μg/mouse, Bioxell) was delivered to mice intraperitoneally at indicated time points. CD8^+^ T cell depletion was confirmed by flow cytometry analysis 24 to 48 h after depletion.

### Viral burden quantification

For *in vivo* experiments, mouse lungs were collected 5 days after SeV challenge. RNA from mouse lungs was extracted using the Qiagen RNeasy kit. 500 ng of total RNA was converted to cDNA using Biorad iScript cDNA conversion kit. Viral burden was determined by reverse transcription quantitative PCR (RT-qPCR) of the Sendai Matrix (M) protein normalized to house-keeping gene 18SRNA. 18S forward primer – GTAACCCGTTGAACCCCATT; reverse primer – CCATCCAATCGGTAGTAGCG. SeV M gene forward primer – ACTGGGACCCTATCTAAGACAT; reverse primer – TAGTAGCGGAAATCACGAGG. The Limit of quantification (LOQ) was established for the SeV qPCR assay as the highest dilution of the template still maintaining the linearity of the assay. The threshold cycle (CT) value of the LOQ was set as the lower limit for the assay.

### ROS inhibition *in vitro* and *in vivo*

NADPH oxidase activity was inhibited by exposing the cells to GKT137831 (10 μM; Selleckchem) 12 h prior to treatment with Pam2-ODN or PBS. Mitochondrial ROS production was inhibited using the combination of FCCP (400 nM, Cayman Chemicals) and TTFA (200 μM, Cayman Chemicals) for 1 h before Pam2-ODN or PBS treatment. For *in vivo* experiments, mice were aerosolized with 10 ml TTFA (200 mM) and FCCP (800 μM) 2 h before Pam2-ODN aerosolization and 2 h before SeV challenge and then again 4 days after SeV challenge^16^.

### Viral attachment assays

MLE-15 cells were cultured in 24 well plates or chamber slides for treatment with Pam2-ODN and SeV inoculations. Cells were placed on ice 30 min before inoculation with SeV to prevent viral entry into the cells. 4 h after inoculation on ice, cells were vigorously washed 5X with media to remove unattached virus. Cells were then harvested to measure SeV burden using immunofluorescence or flow cytometry. For RT-qPCR assays, epithelial cells were treated with Pam2-ODN or PBS, followed by SeV infection on ice (to prevent viral entry into cells). Virus particles were allowed to attach to the epithelial targets for 4 h on ice. These cells were then extensively washed to remove unattached virus particles, and then the cells were lysed by passing through a syringe 10X. The liberated virus particles were then transferred to naïve epithelial cells that had no prior exposure to Pam2-ODN. SeV M gene expression was assessed by qPCR after 24 h of SeV replication in the new cells. The experimental design is illustrated in Fig 5e. In some experiments, mitoROS inhibitors (FFCP-TTFA) were used before Pam2-ODN treatment to determine the role of Pam2-ODN induced ROS in SeV inactivation prior to internalization.

### Statistics

All statistical analysis was performed using GraphPad Prism software (Version 8 for Windows, GraphPad Software, La Jolla, CA). To determine pairwise differences in viral burden or cell numbers, Student’s *t* test was used. Mouse survival analysis of viral challenges were analyzed using Mantel-Cox test. One-way analysis of variance (ANOVA) with multiple comparisons was used to determined differences between multiple experimental conditions.

## Supplementary materials

**Supplementary Figure 1.**
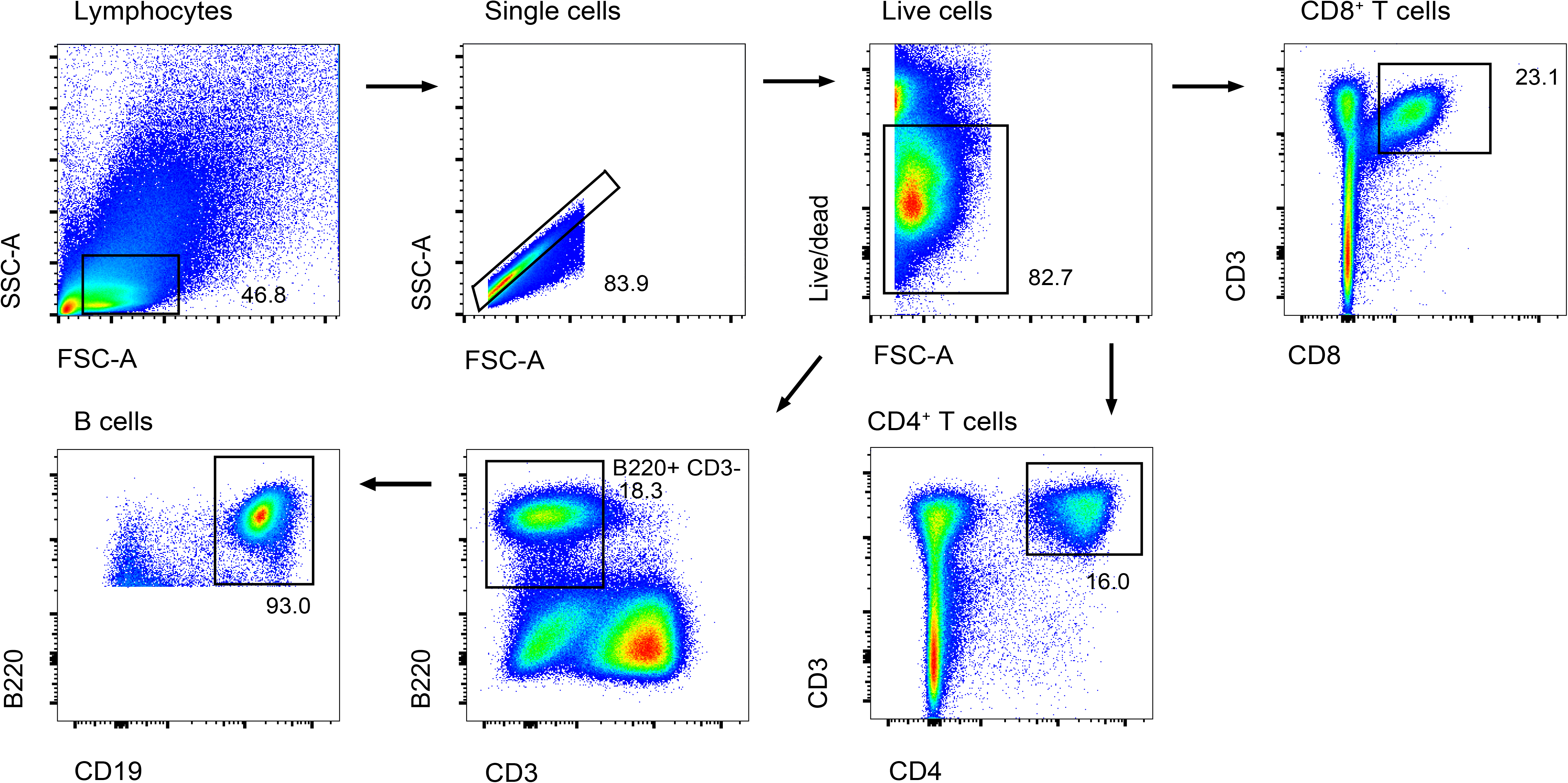
Gating strategy for flow cytometry of lung T and B cells.

**Supplementary Figure 2.**
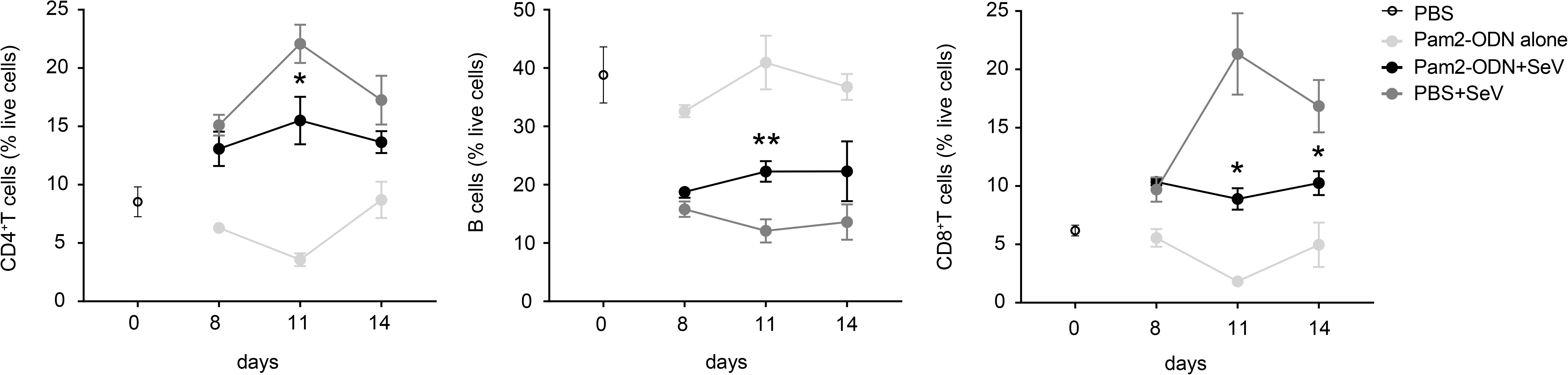
Pam2-ODN pretreatment reduced SeV induced lymphocytes. Disaggregated mouse lung cells positive for CD4^+^ T cells, CD19^+^ B220^+^ B cells, CD8^+^ T cells assessed by flow cytometry in perfused lungs from mice treated with PBS or Pam2-ODN on various days of SeV challenge. \**p*<0.05 compared to PBS+SeV, \*\**p*<0.01 compared to PBS+SeV.

**Supplementary Figure 3.**
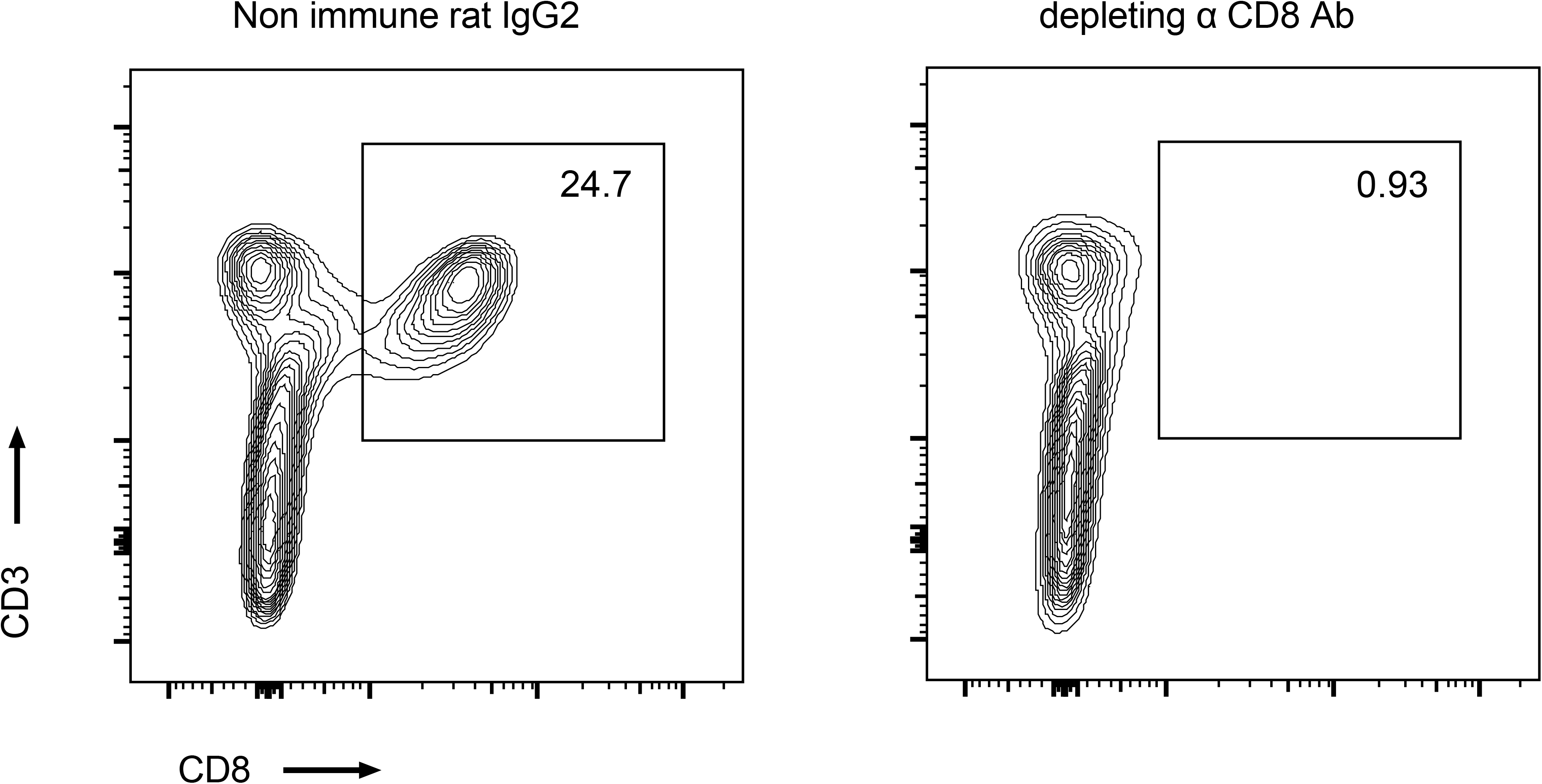
CD8^+^ T cell depleting antibody reduced lung CD8^+^ T cells.

**Supplementary Figure 4.**
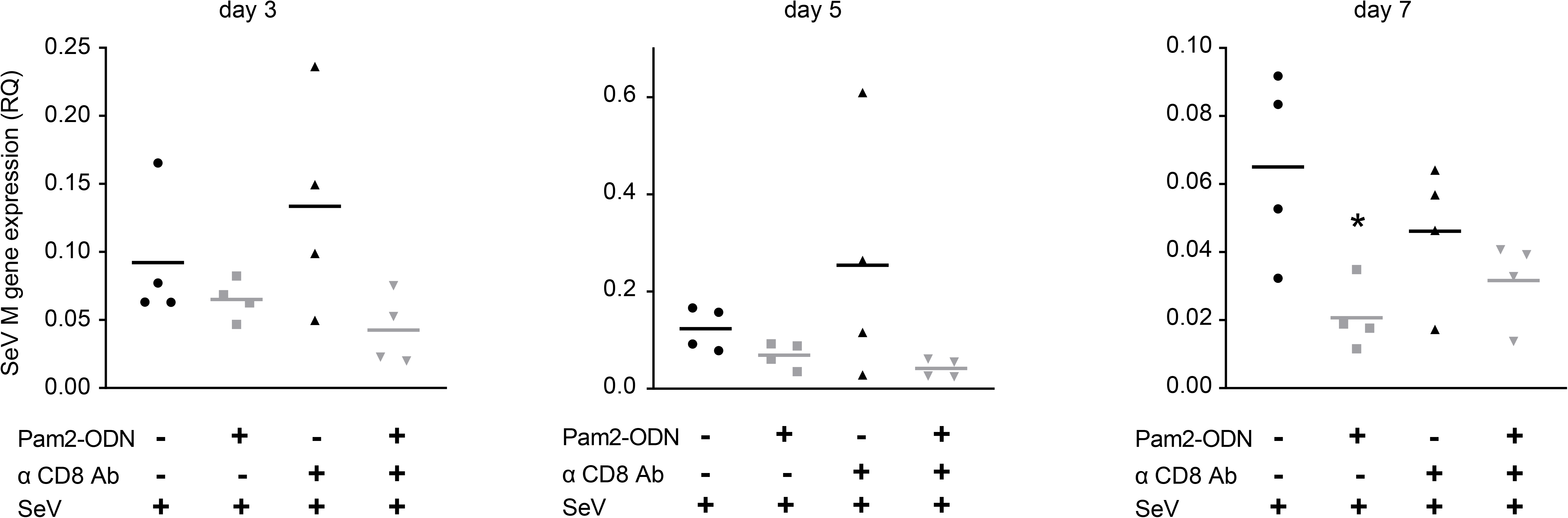
Lung SeV burden 3, 5 and 7 days after SeV challenge in mice treated with PBS or Pam2-ODN with or without CD8^+^ T cell depletion. \**p*<0.05 compared to PBS treated group.

## Author contributions

S.W. designed and performed the experiments, analyzed the data, and wrote the manuscript. J.R.F., A.M.J., D.L.G. and J.P.G performed experiments. M.J.T., B.F.D. conceptualized the project and critically reviewed the data. S.E.E. conceptualized the project, designed experiments, provided critical evaluation of data and edited the manuscript.

## Acknowledgments

The authors would like to thank Dr. Yongxing Wang for optimization of ROS inhibition experiments. M.J.T., B.F.D., and S.E.E. are authors on U.S. patent 8,883,174, “Stimulation of Innate Resistance of the Lungs to Infection with Synthetic Ligands.” M.J.T., B.F.D., and S.E.E. own stock in Pulmotect, Inc., which holds the commercial options on these patent disclosures. All other authors declare that no conflict of interest exists. This study was supported by NIH grants R01 HL117976, DP2 HL123229 and R35 HL144805 to S.E.E.

